# Soluble amyloid-β precursor peptide does not regulate GABA_B_ receptor activity

**DOI:** 10.1101/2022.08.02.502499

**Authors:** Pascal Dominic Rem, Vita Sereikaite, Diego Fernandez-Fernandez, Sebastian Reinartz, Daniel Ulrich, Thorsten Fritzius, Luca Trovò, Salome Roux, Ziyang Chen, Philippe Rondard, Jean-Philippe Pin, Jochen Schwenk, Bernd Fakler, Martin Gassmann, Tania R. Barkat, Kristian Strømgaard, Bernhard Bettler

**Author notes:** Correspondence should be addressed to B.B.

## Abstract

Amyloid-β precursor protein (APP) regulates neuronal activity through the release of secreted APP (sAPP) acting at cell-surface receptors. APP and sAPP were reported to bind to the extracellular sushi domain 1 (SD1) of GABA_B_ receptors (GBRs). A 17 amino-acid peptide (APP17) derived from APP was sufficient for SD1 binding and shown to mimic the inhibitory effect of sAPP on neurotransmitter release and neuronal activity. The functional effects of APP17 and sAPP were similar to those of the GBR agonist baclofen and blocked by a GBR antagonist. These experiments led to the proposal that sAPP activates GBRs to exert its neuronal effects. However, whether APP17 and sAPP indeed influence classical GBR signaling pathways in heterologous cells was not analyzed. Here, we confirm that APP17 binds to GBRs with nanomolar affinity. However, biochemical and electrophysiological assays indicate that APP17 does not influence GBR activity in heterologous cells. Moreover, we found no evidence for APP17 regulating K^+^ currents in cultured neurons, neurotransmitter release in brain slices, or neuronal activity *in vivo*. Our results show that APP17 is not a functional GBR ligand and indicate that sAPP exerts neuronal effects through receptors other than GBRs.

## Introduction

Amyloid precursor protein (APP or A4 protein) is a transmembrane protein that undergoes proteolytic processing by secretases. The amyloidogenic pathway generates amyloid-β peptides (Aβ) that are key etiological agents of Alzheimer’s disease (AD). The competing non-amyloidogenic pathway generates secreted APP (sAPP) variants that modulate spine density, synaptic transmission, plasticity processes, and rescue synaptic deficits in *APP*^-/-^ mice (Muller et al., 2017; Haass and Willem, 2019; Tang, 2019). It is assumed that cell surface receptors mediate the synaptic effects of sAPP (Richter et al., 2018; Haass and Willem, 2019; Tang, 2019; Barthet and Mulle, 2020). It was recently proposed that sAPP acts at G protein-coupled GABA_B_ receptors (GBRs) to modulate synaptic transmission and neuronal activity (Rice et al., 2019)(reviewed by (Haass and Willem, 2019; Korte, 2019; Tang, 2019; Yates, 2019; Barthet and Mulle, 2020)). GBRs are attractive candidates for mediating the functional effects of sAPP because they regulate neurotransmitter release, neuronal inhibition, and synaptic plasticity processes by reducing cAMP levels and gating Ca^2+^ and K^+^ channels (Luscher and Slesinger, 2010; Gassmann and Bettler, 2012; Pin and Bettler, 2016; Barthet and Mulle, 2020).

GBRs are composed of GB1a or GB1b subunits with a GB2 subunit, which generates GB1a/2 and GB1b/2 receptors (Pin and Bettler, 2016). GB1a differs from GB1b by the presence of two N-terminal sushi domains (SD1/2). GB1a/2 and GB1b/2 receptors predominantly localize to pre- and postsynaptic sites, respectively (Vigot et al., 2006). APP and sAPP bind to SD1 of GB1a (Schwenk et al., 2016; Dinamarca et al., 2019; Rice et al., 2019). Synthetic APP peptides of 9 or 17 amino acid residues, termed APP9 and APP17, are sufficient for binding and inducing a stable conformation in SD1 (Rice et al., 2019; Feng et al., 2021; Yang et al., 2022). APP, sAPP and APP17 bound to recombinant SD1 with a *K_D_* of 183, 431 and 810 nM, respectively (Dinamarca et al., 2019; Rice et al., 2019). Consistent with an action on presynaptic GB1a/2 receptors, sAPP and APP17 reduced the frequency of miniature excitatory postsynaptic currents (mEPSCs) in brain slices, similar to the orthosteric GBR agonist baclofen (Rice et al., 2019). The antagonist CGP55845 reduced the inhibitory effect of APP17 on the mEPSC frequency, further supporting that APP17 activates GBRs (Rice et al., 2019). Moreover, APP17 inhibited neuronal activity in the hippocampus of anesthetized mice (Rice et al., 2019), consistent with a GBR-mediated inhibition of glutamate release and/or activation of postsynaptic K^+^ currents. Based on these experiments, Rice and colleagues proposed that APP17 and sAPP are functional GBR ligands. However, no evidence was presented that sAPP or APP17 regulates classical GBR-activated G protein signaling pathways, which is necessary to establish a direct action at GBRs. In separate studies, the binding of APP to GB1a/2 receptors *in cis* was shown to mediate receptor transport to presynaptic sites and to stabilize receptors at the cell surface (Hannan et al., 2012; Dinamarca et al., 2019). Accordingly, *APP*^-/-^ mice exhibited a 75% decrease of axonal GBRs in hippocampal neurons, which significantly reduced GBR-mediated presynaptic inhibition (Dinamarca et al., 2019), as already observed earlier (Seabrook et al., 1999). However, sAPP had no effect on GBR-mediated G protein activation in transfected HEK293 cells (Dinamarca et al., 2019). Thus, it remains controversial whether sAPP and APP17 are functional ligands at GBRs or not.

The reported effects of APP17 on synaptic release and neuronal activity (Rice et al., 2019) suggest that APP17 acts as a positive allosteric modulator (PAM) or ago-PAM (PAM with agonistic properties) at GBRs. However, in principle, APP17 could also increase constitutive activity of GBRs (Grunewald et al., 2002; Rajalu et al., 2015) by binding to SD1 and/or displacing APP from SD1. To clarify whether APP17 influences GBR activity, we studied the effects of APP17 on classical GBR signaling pathways in transfected HEK293T cells, cultured neurons, acute hippocampal slices and in anesthetized mice. Our experiments confirm that APP17 binds with nanomolar affinity to purified recombinant SD1/2 protein and to GB1a/2 receptors expressed in HEK293T cells. However, in our hands, APP17 neither induced conformational changes consistent with GBR activation nor influenced GBR-mediated G protein activity, cAMP inhibition or Kir3-type K^+^ currents. APP17 also failed to modulate constitutive GBR activity in the absence or presence of APP expressed *in cis* or *in trans*. Moreover, APP17 neither influenced K^+^ currents in cultured hippocampal neurons, nor reduced the amplitude of evoked EPSCs in acute hippocampal slices or modulated neuronal activity in living mice. Thus, our *in vitro* and *in vivo* data indicate that receptors other than GBRs mediate the synaptic effects of sAPP.

## Results

### APP17 binds to purified recombinant SD1/2 protein and GB1a/2 receptors expressed in HEK293T cells

We purchased APP17 and scrambled sc-APP17 peptides from the same commercial provider as Rice and colleagues (Rice et al., 2019) (Figure 1A). For displacement experiments, we additionally synthesized fluorescent APP17-TMR and sc-APP17-TMR peptides labeled with TAMRA (5(6)-carboxytetramethylrhodamine, Figure 1A). ESI-LC-MS and RP-UPLC analysis revealed that all peptides had the expected molecular weight and purity (Figure 1A). Isothermal titration calorimetry (ITC) showed that APP17 interacts with purified recombinant SD1/2 protein (Schwenk et al., 2016) in solution with a ~1:1 stoichiometry and a K*_D_* of 543 nM (Figure 1B). This agrees well with the published K*_D_* of 810 nM for binding of APP17 to SD1 (Rice et al., 2019). In contrast, sc-APP17 showed no detectable binding (K_D_ > 300 μM). APP17-TMR exhibited significantly more binding to HEK293T cells expressing GB1a/2 receptors than sc-APP17-TMR (Figure 1C). Accordingly, 10 μM APP17 but not sc-APP17 displaced 1 μM APP17-TMR from GB1a/2 receptors (Figure 1C). In all subsequent experiments, we used the commercial APP17 peptide validated for binding to recombinant SD1/2 protein and GB1a/2 receptors expressed in HEK293T cells.

**Figure 1.**
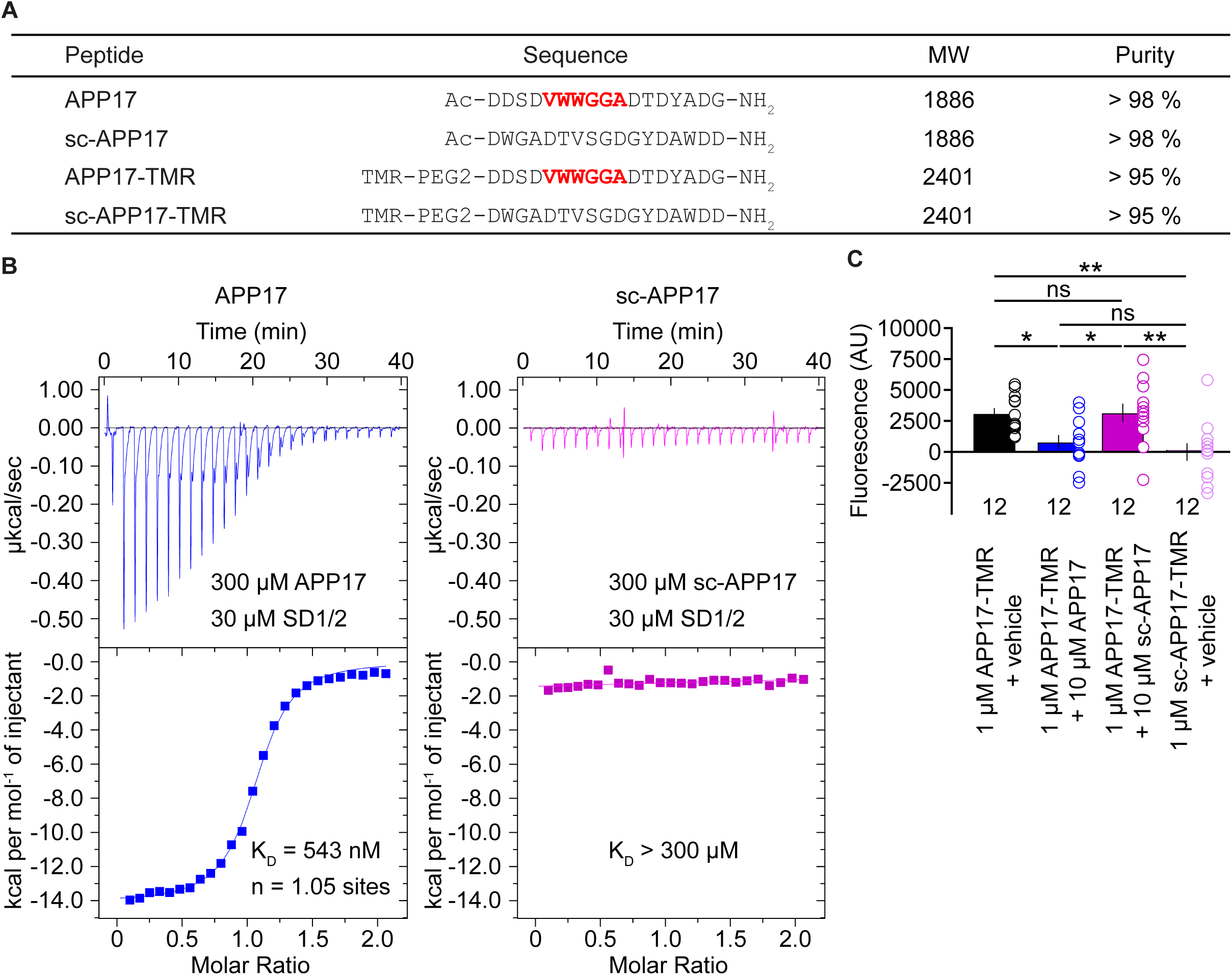
Characterization of synthetic APP17 and sc-APP17 peptides. (**A**) Sequence alignment of APP17, sc-APP17, APP17-TMR and sc-APP17-TMR peptides. Residues critical for SD1 binding are shown in red. (**B**) Representative ITC diagrams of the titrations of SD1/2 protein in solution (30 μm) with APP17 (blue) or sc-APP17 (magenta) (300 μm in the syringe); raw heat signature (top) and integrated molar heat release (bottom). The calculated stoichiometry of APP17:SD1/2 protein is 1.05, the K_D_ 543 nM. sc-APP17 showed no binding to SD1/2 protein. (**C**) Bar graph showing APP17-TMR (1 μM) binding to GB1a/2 receptors in HEK293T cells in the presence of vehicle (black), 10 μM APP17 (blue), and 10 μM sc-APP17 (magenta). sc-APP17-TMR (1 μM) served as a negative control. The background fluorescence of sc-APP17-TMR (1 μM) at HEK293T cells transfected with empty vector was subtracted. Data are means ± SEM. The number of independent experiments is indicated. ns = not significant, *p < 0.05, **p < 0.01, one-way ANOVA with Holm-Sidak’s multiple comparisons test. Source file containing ICT and TMR fluorescence data is available in Figure 1 – source data 1.

### APP17 does not induce the active state of GB1a/2 receptors

Upon binding of APP17, SD1 adopts a stable conformation (Rice et al., 2019) that possibly influences GBR activity allosterically. We used a fluorescence resonance energy transfer (FRET)-based conformational sensor in transfected HEK293 cells to analyze whether APP17 induces the inter-subunit rearrangement associated with GBR activation (Lecat-Guillet et al., 2017). The FRET sensor is based on GB1a and GB2 subunits fused with ACP and SNAP (Figure 2A), respectively. These tags are then enzymatically modified with time-resolved FRET compatible fluorophores (HA-GB1a-ACP with CoA-Lumi4-TB (Donor), Flag-SNAP-GB2 with SNAP-RED (Acceptor)). This FRET sensor discriminates between GBR agonists with different efficacies and between PAMs with distinct modes of action (Lecat-Guillet et al., 2017). For FRET experiments, we used APP17 and sc-APP17 at 1 μM and 10 μM. These concentrations are above the K*_D_* of APP17 and comparable to those used in previous functional experiments (25 nM - 5 μM) (Rice et al., 2019). As expected, GABA decreased FRET in a dose-dependent manner (Figure 2B). APP17 at 10 μM, when applied alone (basal, Figure 2B), significantly increased instead of decreased FRET (Figure 2B,C). This FRET increase can be rationalized in two ways. First, since GBRs exhibit constitutive activity (Grunewald et al., 2002; Rajalu et al., 2015), there is an equilibrium between active and inactive states of the receptor. Constitutive activity decreases basal FRET because a fraction of receptors is in the active state. Ligands stabilizing the inactive conformation are therefore increasing basal FRET. Therefore, APP17 is potentially an inverse agonist of GBRs. However, since all functional experiments reveal no inverse agonistic properties (see below), APP17 binding to SD1 likely influences the positioning of the SNAP-tag located on top of GB2, thereby decreasing the mean distance between the fluorophores and increasing FRET efficacy. Of note, APP17 lacks allosteric properties, as it did not significantly alter GABA potency (Figure 2D). In summary, the FRET conformational sensor provides no evidence for APP17 allosterically promoting receptor activation.

**Figure 2.**
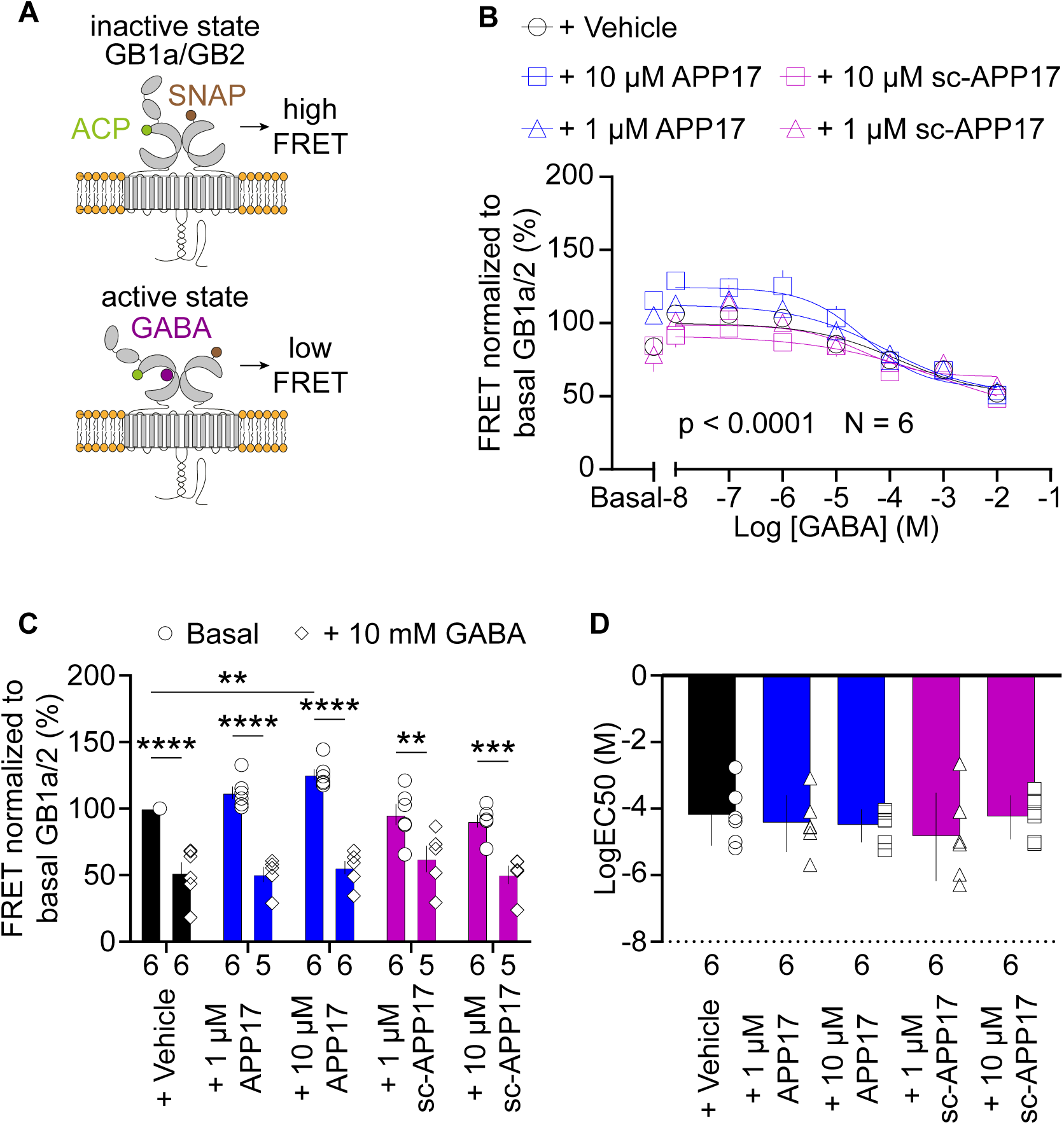
APP17 does not induce the active state of GB1a/2 receptors. (**A**) Assay measuring intersubunit FRET between fluorophore labelled ACP and SNAP tags in the GB1a and GB2 subunits, respectively. In the absence of receptor agonists, the ACP and SNAP tags are in close proximity resulting in high FRET. Activation of GB1a/2 receptors induces a conformational change in the extracellular domains leading to a reduction in FRET. (**B**) GABA dose-response curves in the presence of APP17 (blue) or sc-APP17 (magenta) at 1 μM (triangles) and 10 μM (squares) or vehicle (black) exhibit significant differences. (**C**) Bar graphs showing FRET in the presence of 1 μM or 10 μM APP17 (blue) and sc-APP17 (magenta) or vehicle (black). Under basal conditions (circles), the presence of 10 μM APP17 resulted in a significant increase of FRET, whereas no significant changes in FRET were observed for all other conditions when compared to vehicle. In the presence of 10 mM GABA (diamonds) no significant differences in FRET were detected with APP17 or sc-APP17 at 1 μM or 10 μM compared to vehicle. In all conditions, the presence of 10 mM GABA induced a significant reduction in FRET compared to basal. (**D**) LogEC_50_ values of individual GABA dose-response curves exhibit no significant differences between conditions. Data are means ± SEM. Three outliers were identified in **c** using the ROUT method (PRISM) with Q = 1% (source file). The number of independent experiments is indicated. **p < 0.01, ***p < 0.001, ****p < 0.0001, two-way ANOVA with Sidak’s multiple comparisons test. Source file containing FRET data is available in Figure 2 – source data 1.

### APP17 does not influence GB1a/2 receptor-mediated G protein activation

We directly tested whether APP17 activates or modulates G protein activation in transfected HEK293T cells expressing GBRs. We used a bioluminescent resonance energy transfer (BRET) assay monitoring dissociation of Gα from Gβγ upon receptor activation (Turecek et al., 2014) (Figure 3A). Application of GABA to cells expressing GB1a/2 together with Gαo-RLuc, Venus-Gγ2 and Gβ2 lead to a BRET decrease between Gαo-RLuc and Venus-Gγ2 (Figure 3A). Subsequent blockade of GB1a/2 receptors by the inverse agonist CGP54626 (Grunewald et al., 2002) increased BRET due to re-association of G protein subunits (Figure 3A). Of note, CGP54626 increased BRET above baseline, consistent with substantial constitutive activity of GBRs (Grunewald et al., 2002; Rajalu et al., 2015). Accordingly, application of CGP54626 alone to transfected HEK293T cells also increased BRET (Figure 3A). Application of 10 μM GABA did not overcome receptor inhibition by 10 μM CGP54626 (Figure 3A). The presence of APP695 *in cis* did not alter constitutive activity of the receptor (Figure 3A). APP17 and sc-APP17 at 1 or 10 μM did not significantly influence receptor activity while subsequent GABA application to the same cells induced the expected BRET decrease (Figure 3B). GABA-induced BRET decreases were similar in the presence of APP17 or sc-APP17 (Figure 3B). These experiments indicate that APP17 at 1 or 10 μM exerts no agonistic, inverse agonistic, antagonistic or allosteric properties at GB1a/2 receptors. Moreover, application of APP17 or sc-APP17 to HEK293T cells expressing GB1a/2 receptors and APP695 *in cis* or *in trans* had no effect on GBR activity (Figure 3C,D). Subsequent application of GABA was equally effective in decreasing BRET in the presence of APP17 or sc-APP17 (Figure 3C,D). These experiments show that APP17 does not modulate GBR-mediated G protein activation in the absence or presence of APP695.

**Figure 3.**
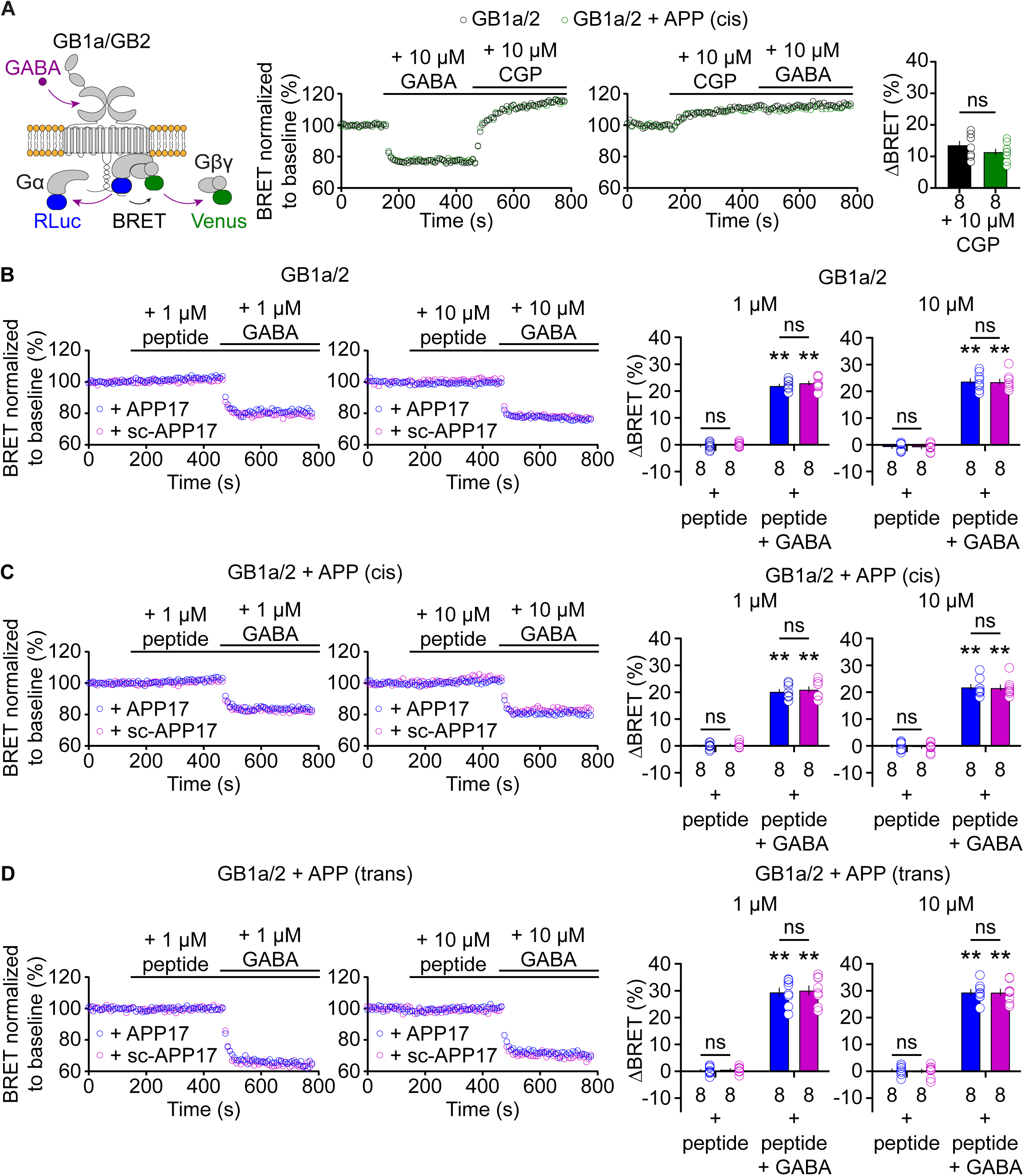
APP17 is not an agonist, inverse agonist, antagonist or allosteric modulator at GB1a/2 receptors expressed in HEK293T cells in a BRET assay monitoring G protein activation. (**A**) *Left* Assay measuring BRET between Gα_o_-RLuc and Venus-Gγ_2_. GB1a/2 receptor activation leads to dissociation of the heterotrimeric G protein and a consequent decrease in BRET. *Right* Individual experiments showing GABA-induced BRET changes at GB1a/2 receptors. The inverse agonist CGP54626 reverses GABA-induced BRET changes above baseline, indicating constitutive GB1a/2 receptor activity. Likewise, direct application of CGP54626 increased BRET levels above baseline. Subsequent application of GABA did not overcome receptor inhibition. Bar graphs summarize CGP54626-induced BRET changes. Note that application of CGP54626 resulted in similar inhibition of constitutive GB1a/2 receptor activity in the absence (black) or presence (green) of APP695 *in cis*. (**B**) Neither APP17 (blue) nor sc-APP17 (magenta) at 1 μM (left) or 10 μM (right) altered BRET in cells expressing GB1a/2 receptors. In the same cells, GABA at 1 μM (left) and 10 μM (right) induced the expected decrease in BRET. The GABA-induced BRET change is similar in the presence of APP17 and scAPP17, indicating the absence of allosteric properties of the peptides at GB1a/2 receptors. Bar graphs summarize BRET changes determined in experiments as shown to the left. (**C**,**D**) Neither APP17 (blue) nor sc-APP17 (magenta) at 1 μM (left) or 10 μM (right) altered BRET in cells expressing GB1a/2 receptors together with APP695 *in cis* (C) or *in trans* (D). Bar graphs summarize BRET changes. Data are means ± SEM. The number of independent experiments is indicated in the bar graphs. ns = not significant, Two-way ANOVA with Sidak’s multiple comparisons test. **p < 0.01, One sample Wilcoxon test (non-parametric) against 0. Source file containing BRET data is available in Figure 3 – source data 1.

### APP17 does not influence GB1a/2 receptor-mediated Gα signaling

Assays measuring Gαi signaling provide another means to study possible functional effects of APP17 on GBR activity. We analyzed whether APP17 influences GB1a/2-mediated Gαi signaling using an assay monitoring cAMP-dependent Protein Kinase A (PKA) activity in transfected HEK293T cells. This assay is based on regulatory and catalytic PKA subunits tagged with the N- or C-terminal fragments of RLuc (R-RLuc-N, C-RLuc-C) (Stefan et al., 2007). GB1a/2 receptor activation by 10 μM GABA inhibits adenylyl cyclase, which inactivates PKA and increases luminescence due to association of R-RLuc-N with C-RLuc-C (Figure 4A). Blockade of GB1a/2 receptors with 10 μM CGP54626 decreased luminescence below baseline, again revealing substantial constitutive receptor activity in this assay system (Figure 4A). APP17 or sc-APP17 at 10 μM exhibited no agonistic, inverse agonistic or antagonistic properties at GB1a/2 receptors in the PKA assay (Figure 4B). GABA-mediated PKA inactivation was comparable in the presence of APP17 or sc-APP17, again supporting that APP17 does not act as a PAM (Figure 4B). Moreover, APP17 or sc-APP17 did not significantly alter GB1/2 receptor activity in the presence of APP695 *in cis* (Figure 4C).

**Figure 4.**
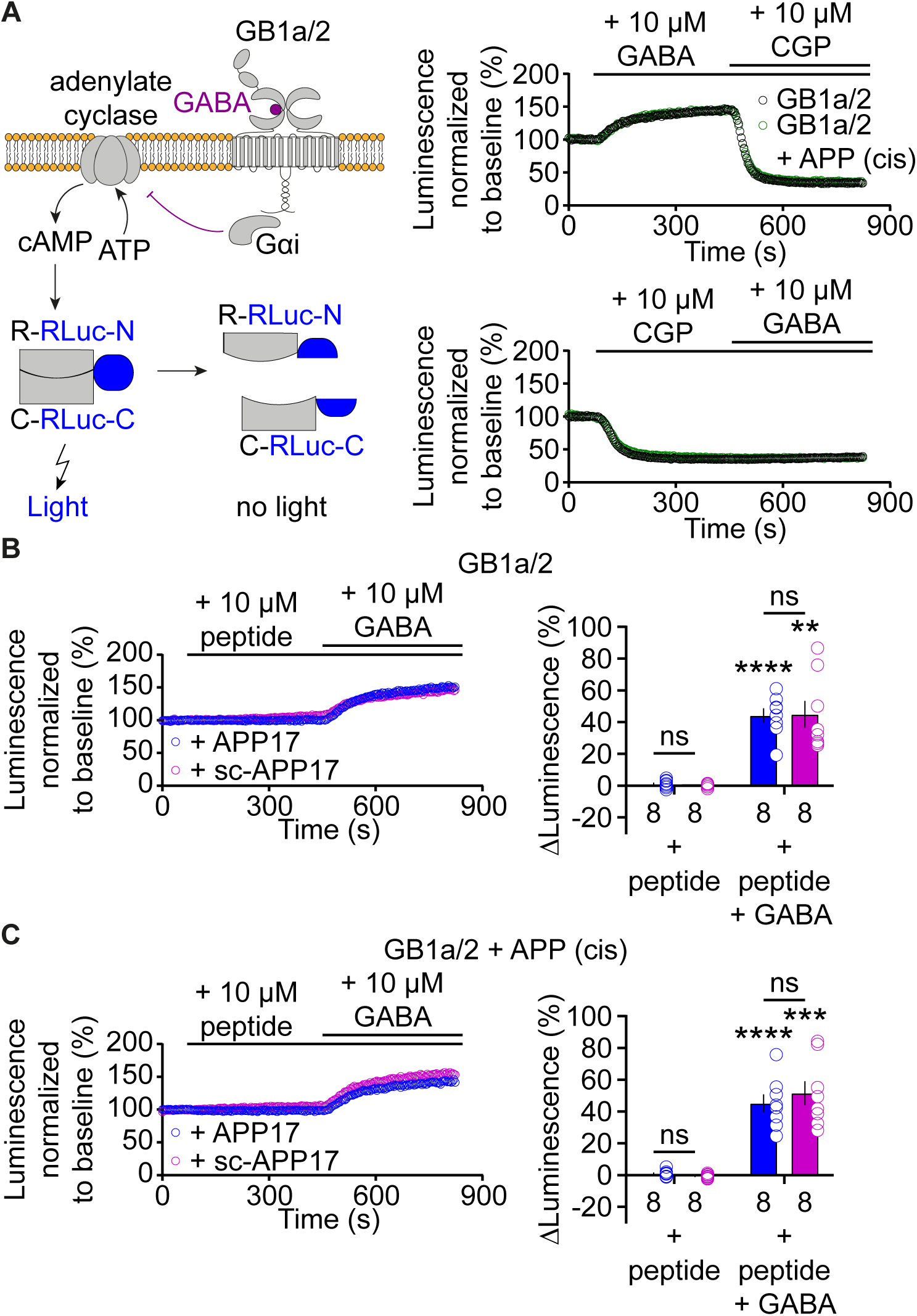
APP17 is not an agonist, inverse agonist, antagonist or allosteric modulator at GB1a/2 receptors expressed in HEK293T cells in an assay monitoring Gα_i_ signaling. (**A**) *Left* Assay monitoring dissociation of the regulatory (R) and catalytic (C) subunits of the tetrameric PKA holoenzyme upon cAMP binding. PKA subunits were tagged with N- or C-terminal fragments of RLuc (R-RLuc-N, C-RLuc-C). GB1a/2 receptor activation by GABA reduces intracellular cAMP levels, promotes reconstitution of RLuc activity and increases luminescence. *Right* Individual experiments showing GABA-induced luminescence changes. Blockade of GB1a/2 receptor activity with CGP54626 decreased luminescence below baseline, indicating constitutive GB1a/2 receptor activity. Subsequent application of GABA did not overcome receptor inhibition. (**B**) Neither APP17 (blue) nor sc-APP17 (magenta) altered luminescence in HEK293T cells expressing GB1a/2 receptors. In the same cells, GABA induced the expected luminescence increases. GABA-induced luminescence changes are similar in the presence of APP17 and sc-APP17. Bar graphs summarize the luminescence changes. (**C**) Neither APP17 (blue) nor sc-APP17 (magenta) induced luminescence changes in HEK293T cells expressing GB1a/2 receptors together with APP695 *in cis*. Application of GABA to the same cells resulted in the expected luminescence increases. GABA-induced luminescence changes are similar in the presence of APP17 or sc-APP17. Bar graphs summarize the luminescence changes. Data are means ± SEM. The number of independent experiments is indicated. ns = not significant, Two-way ANOVA with Sidak’s multiple comparison test. **p < 0.01, ***p < 0.001; ****p < 0.0001, One sample t-test against 0. Source file containing PKA luminescence data is available in Figure 4 – source data 1.

It is conceivable that the APP17 concentrations used are not optimal for detecting functional effects at recombinant GB1a/2 receptors. Therefore, we determined APP17 dose-response curves using an accumulation assay based on artificially coupling GB1a/2 receptors to phospholipase C (PLC) via chimeric Gα_qi_ (Conklin et al., 1993) (Figure 5A). PLC activity was monitored with a serum responsive element-luciferase (SRE-Luciferase) reporter amplifying the receptor response (Yoo et al., 2017). Increasing concentrations of GABA yielded similar sigmoidal dose-response curves in the absence and presence of APP695 expressed *in cis* or *in trans* (Figure 5 – figure supplement 1), showing that binding of APP695 does not influence receptor activity. APP17 or sc-APP17 lacked agonistic properties at concentrations up to 100 μM, the highest concentration tested (Figure 5A). CGP54626 blocked constitutive and GABA-induced receptor activity (Figure 5B). APP17 or sc-APP17 at concentrations of 1 and 10 μM did not influence constitutive GB1a/2 receptor activity (Figure 5B). Pre-incubation with 1 µM or 10 µM of APP17 or sc-APP17 did not significantly influence the GABA dose-response curve in the absence (Figure 5C) or presence of APP695 expressed *in cis* (Figure 5D), corroborating that the peptides lack agonistic, PAM, inverse agonistic or antagonistic properties.

**Figure 5.**
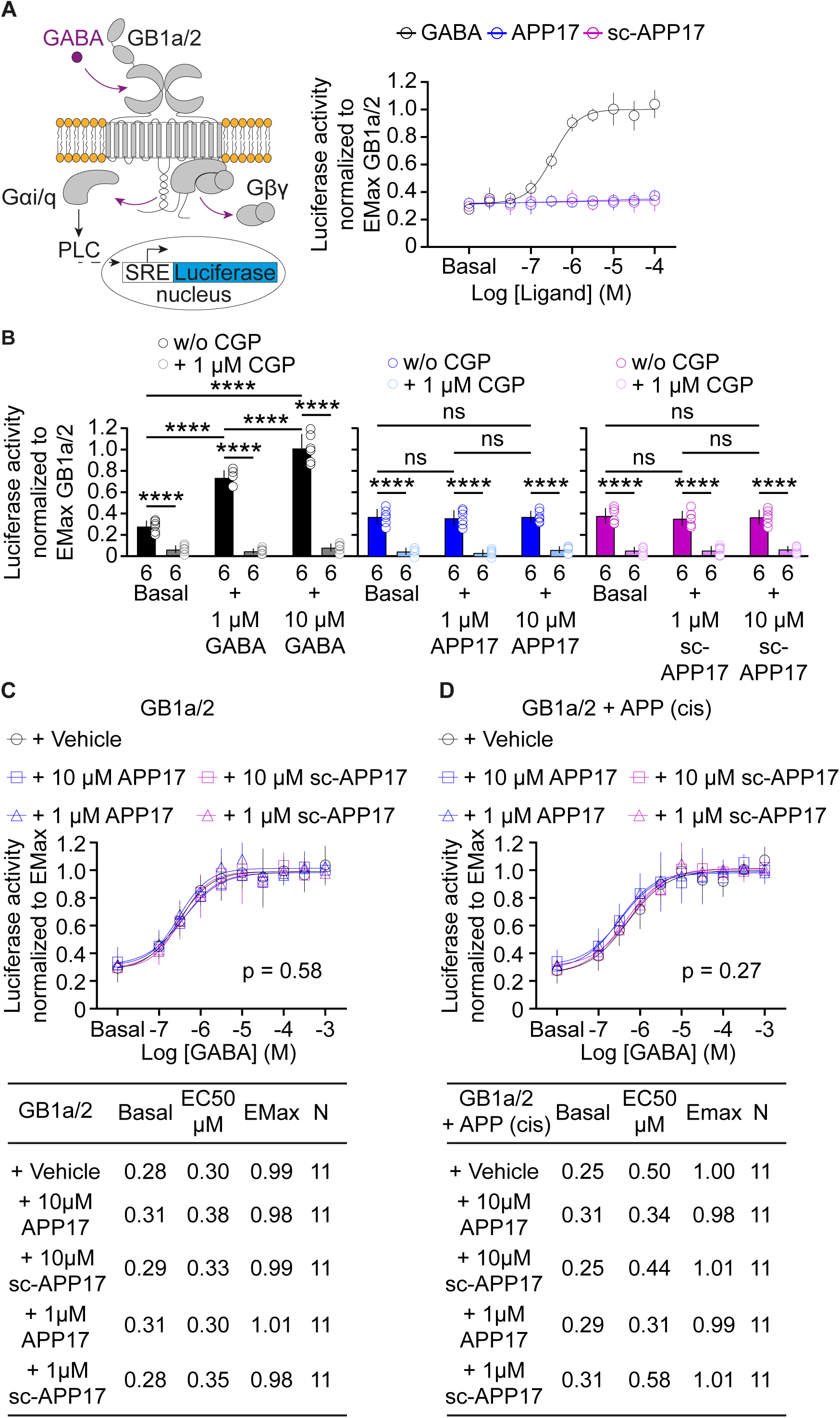
APP17 is not an agonist, inverse agonist, allosteric modulator or antagonist at GB1a/2 receptors expressed in HEK293T cells when monitoring Ga_qi_ signaling in an accumulation assay. (**A**) *Left* Assay monitoring PLC dependent FLuc expression under control of the serum response element (SRE). GB1a/2 receptors were artificially coupled to PLC by stably expressing the chimeric G protein subunit Gα_qi_. GB1a/2 receptors and SRE-FLuc reporter were transiently expressed in HEK293T-Gα_qi_ cells. *Right* Dose-response curve showing that GABA (black) but not APP17 (blue) or sc-APP17 (magenta) induces FLuc activity in transfected cells. (**B**) CGP54626 blocked constitutive and GABA-induced FLuc activity in transfected cells. Constitutive GB1a/2 receptor activity is unchanged in the presence of APP17 (middle) or sc-APP17 (right) at 1 or 10 μM, indicating the absence of inverse agonistic properties of the peptides at GB1a/2 receptors. (**C**,**D**) APP17 (blue) or sc-APP17 (magenta) at 1 μM (triangles) or 10 μM (squares) did not significantly alter GABA dose-response curves in the absence (C) or presence (D) of APP695 *in cis*, indicating that the peptides do not allosterically regulate GB1a/2 receptors. Tables show basal, EC_50_ and Emax values derived from the curve fits. All data are mean ± SD. The number of independent experiments is indicated in the bar graphs or tables. Linear regression curve fit of 6 (APP17, sc-APP17, A) independent experiments per condition. Non-linear regression curve fits of 6 (GABA, A) or 11 (C,D) independent experiments per condition. p = 0.58, p = 0.27, extra sum-of-squares F test. Source file containing FLuc activity data is available in Figure 5 – source data 1.

### APP17 peptide does not influence [^35^S]GTPγS binding in brain tissue

Native GBRs form multi-protein complexes with auxiliary proteins (Pin and Bettler, 2016; Schwenk et al., 2016). It is conceivable that the functional APP17 effects observed in neurons (Rice et al., 2019) depend on the presence of GBR-associated proteins other than APP that are absent in heterologous expression systems. Binding of the non-hydrolyzable GTP analog [^35^S]guanosine-5’-O-(3-thio)triphosphate ([^35^S]GTPγS) to Gαi/o in brain membranes allows to quantify G protein activation by native GBRs (Galvez et al., 2000; Schuler et al., 2001). GABA dose-response curves for native GBRs in the absence and presence of 1 μM APP17 did not significantly differ from each other and exhibited similar EC_50_ and E_max_ values (Figure 6). This finding supports that 1 μM APP17 has no agonistic, inverse agonistic, antagonistic or allosteric effects at native GBRs.

**Figure 6.**
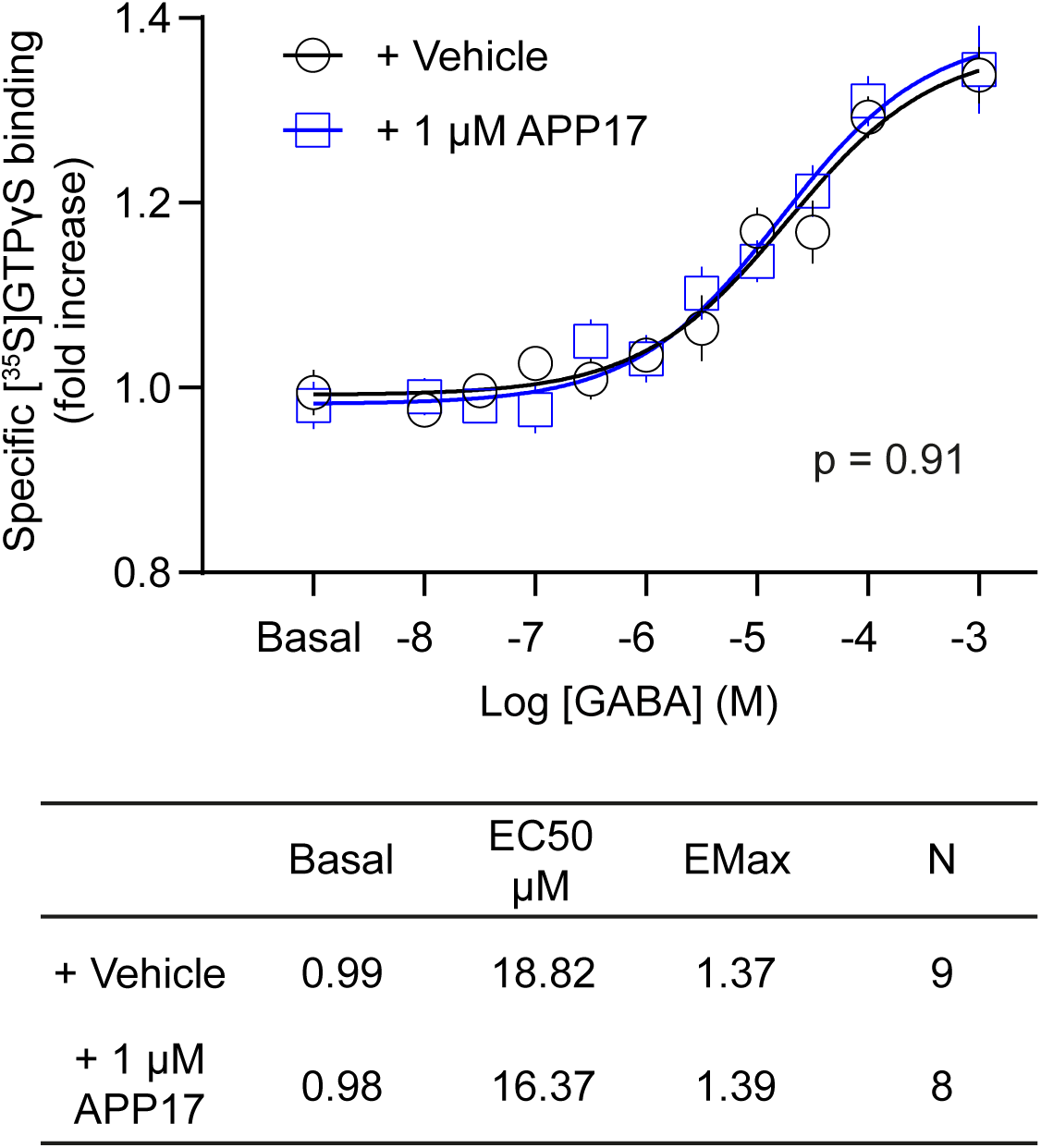
APP17 is not an agonist, antagonist or allosteric modulator at native GB1a/2 receptors in [^35^S]GTPγS binding experiments. [^35^S]GTPγS binding in membrane preparations of WT mice induced by increasing concentrations of GABA is not altered in the presence of APP17 (blue). The table shows basal, EC_50_ and Emax values derived from non-linear regression curve fits. Experiments with vehicle and APP17 were performed with membrane preparations from the same mouse. Data are mean ± SEM. Non-linear regression curve fit of 9 (vehicle) and 8 (APP17) independent experiments with 9 different mice. p = 0.91, extra sum-of-squares F test. Source file containing [^35^S]GTPγS data is available in Figure 6 – source data 1.

### APP17 does not influence GBR-activated K^+^ currents in neurons and transfected HEK293T cells

APP17 signaling through native GBRs may depend on protein-protein interactions that are not preserved in the membrane preparations used for the [^35^S]GTPγS binding assay. Therefore, we tested whether APP17 influences GBR-mediated Gβγ signaling to K^+^ channels in cultured hippocampal neurons using patch clamp electrophysiology, which preserves the native environment of receptors (Schuler et al., 2001; Vigot et al., 2006). Application of 5 µM APP17 or sc-APP17 to hippocampal neurons did not elicit any currents, in contrast to the same concentration of baclofen (Figure 7A,B). Co-application of APP17 or sc-APP17 with baclofen elicited currents of similar amplitudes as baclofen alone (Figure 7B,C), indicating that APP17 exerts no allosteric properties. APP17 or sc-APP17 also did not trigger K^+^ currents in transfected HEK293T cells expressing Kir3 channels, nor did the peptides alter K^+^ currents in the presence of 5 µM GABA (Figure 7 – figure supplement 1). These findings support that APP17 has no agonistic, PAM or antagonistic effects at GBR-activated K^+^ currents.

**Figure 7.**
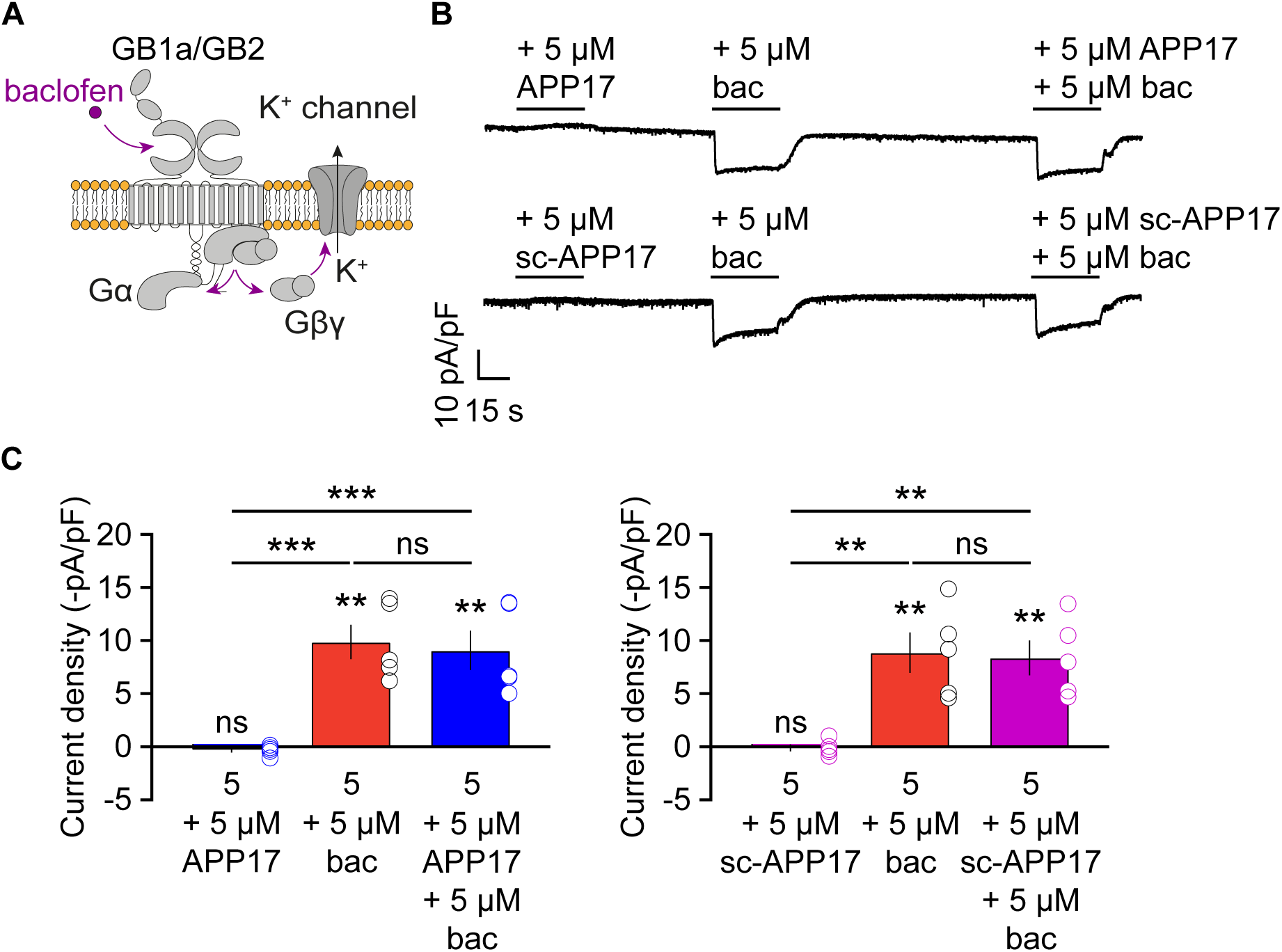
APP17 does not evoke or influence GB1a/2 receptor-mediated K^+^ currents in cultured hippocampal neurons. (**A**) GBR activation with baclofen results in the dissociation of the heterotrimeric G protein and the subsequent activation of K^+^ channels by Gβγ. (**B**) Representative traces showing that neither APP17 (top) nor sc-APP17 (bottom) evoke GB1a/2 receptor-induced K^+^ currents in cultured hippocampal neurons. Application of baclofen alone or in the presence of APP17 or sc-APP17 yielded similar current amplitudes, showing that APP17 does not allosterically modulate baclofen-induced currents. (**C**) Bar graphs showing K^+^ current densities determined in experiments as shown to the top. Data are means ± SEM. The number of independent experiments is indicated in the bar graphs. ns = not significant, **p < 0.01, ***p < 0.001 Paired one-way ANOVA with Holm-Sidak’s multiple comparisons test (to compare different means) and One sample t-test against 0. Source file containing K^+^ current data is available in Figure 7 – source data 1.

### APP17 does not influence evoked EPSC amplitudes in acute hippocampal slices

GB1a/2 receptors are abundant at axon terminals where they inhibit neurotransmitter release (Vigot et al., 2006). Acute exposure of cultured mouse hippocampal neurons to APP17 at 250 nM was shown to inhibit the mEPSC frequency, consistent with an activation of presynaptic GB1a/2 receptors (Rice et al., 2019). A GBR antagonist blocked the effect of APP17 on the mEPSC frequency, supporting a GBR-dependent mechanism. Since all our experiments thus far showed no functional effects of APP17 at GBRs, we next sought to replicate the effects of APP17 at presynaptic GBRs. In our experience, the reduction of the evoked EPSC amplitude provides a better signal-to-noise ratio than the reduction of the mEPSC frequency for assessing presynaptic GBR activity in the hippocampus. Therefore, we studied whether APP17 at 1 μM influences evoked EPSC amplitudes in acute hippocampal slices. Postsynaptic GBR-activated K^+^ currents were blocked with a Cs^+^-based intracellular solution, which allowed to specifically analyze the activity of presynaptic GBRs. Our electrophysiological recordings showed that baclofen but not APP17 was able to reduce the amplitudes of evoked EPSCs (Figure 8).

**Figure 8.**
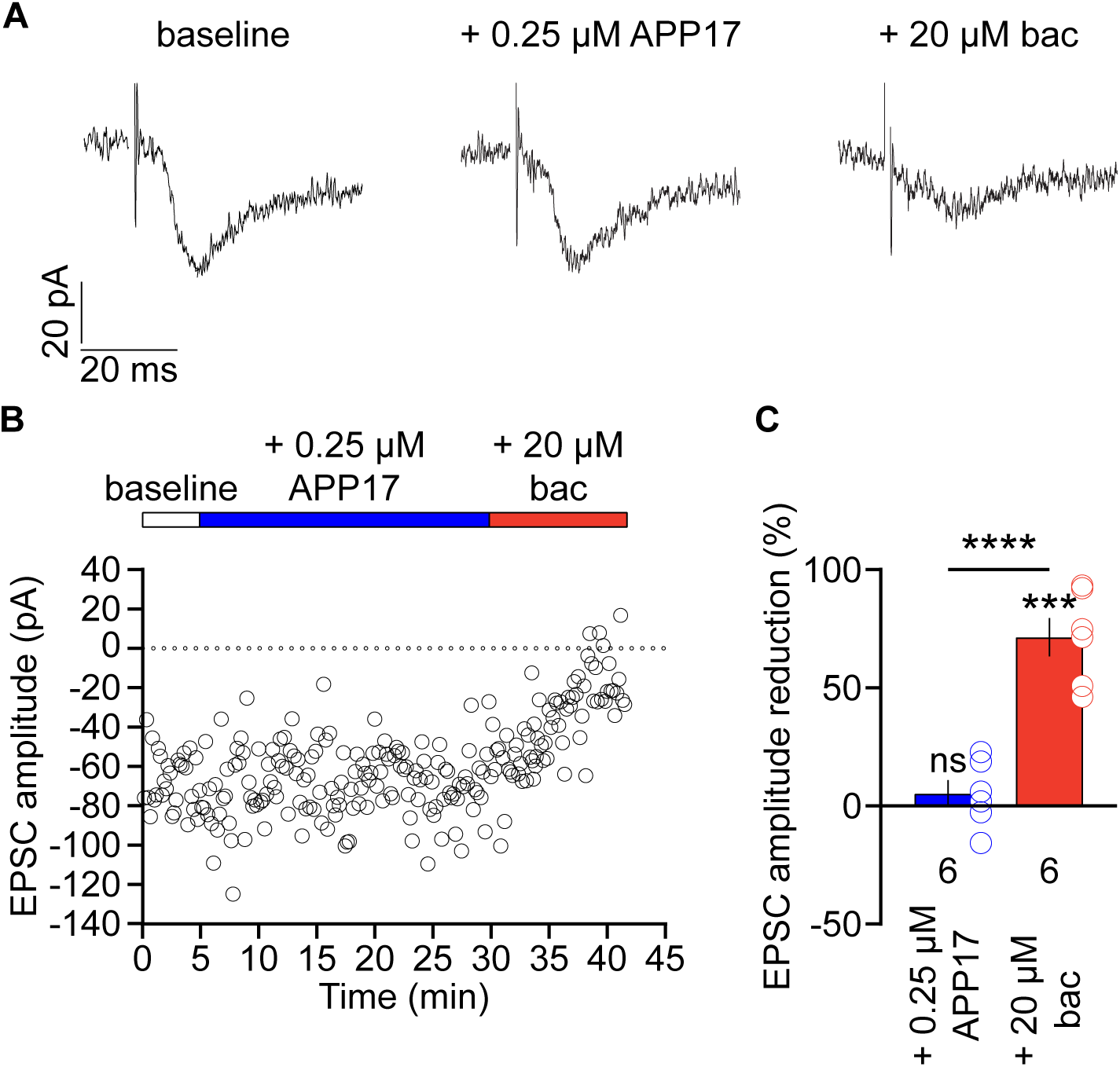
APP17 does not influence the amplitude of evoked EPSCs recorded in CA1 pyramidal neurons of acute hippocampal slices. (**A**) Sample EPSCs at baseline and in the presence of APP17 and baclofen (bac). (**B**) Time course of EPSC amplitudes in a CA1 pyramidal neuron APP17 and baclofen were bath applied as indicated. (**C**) Summary bar graph of the EPSC amplitude reduction in the presence of APP17 and baclofen. The number of recorded neurons from 6 different mice is indicated. ns = not significant; ***p<0.001, one sample t-test against 0; ****p<0.0001, unpaired t-test. Source file containing EPSC data is available in Figure 8 – source data 1.

### APP17 does not influence spontaneous neuronal activity in the auditory cortex of anesthetized mice

Two-photon Ca^2+^ imaging showed that APP17 suppresses neuronal activity of CA1 pyramidal cells in anesthetized mice (Rice et al., 2019). We therefore similarly performed two-photon Ca^2+^ imaging experiments in anesthetized transgenic mice to analyze whether physiological concentrations of APP17 modulate spontaneous activity in cortical neurons, where the density of GBRs in the brain is high (Bischoff et al., 1999). We crossed Ai95(RCL-GCaMP6f)-D mice (Madisen et al., 2015) with Nex-Cre mice (Goebbels et al., 2006) to express the Ca^2+^ indicator GCaMP6f under the Nex-promoter, which allowed us to record Ca^2+^ transients in layer 2/3 neurons of the right auditory cortex. APP17, sc-APP17 and baclofen solutions were perfused over the cortical surface in a fixed sequence (Figure 9A). To control for potential time-dependent changes of spontaneous activity under isoflurane anesthesia (Magnuson et al., 2014), we perfused ACSF before and after perfusion of sc-APP17 and APP17. The results showed that sc-APP17 and APP17 at concentrations of 5 µM had no significant effect compared to ACSF, even after 60 minutes of perfusion (Figure 9D,E, Figure 9 – figure supplement 1). In contrast, 5 µM of baclofen reduced spontaneous Ca^2+^ transients after 15 minutes of perfusion. Therefore, we were unable to confirm that APP17 influences neuronal activity *in vivo*.

**Figure 9.**
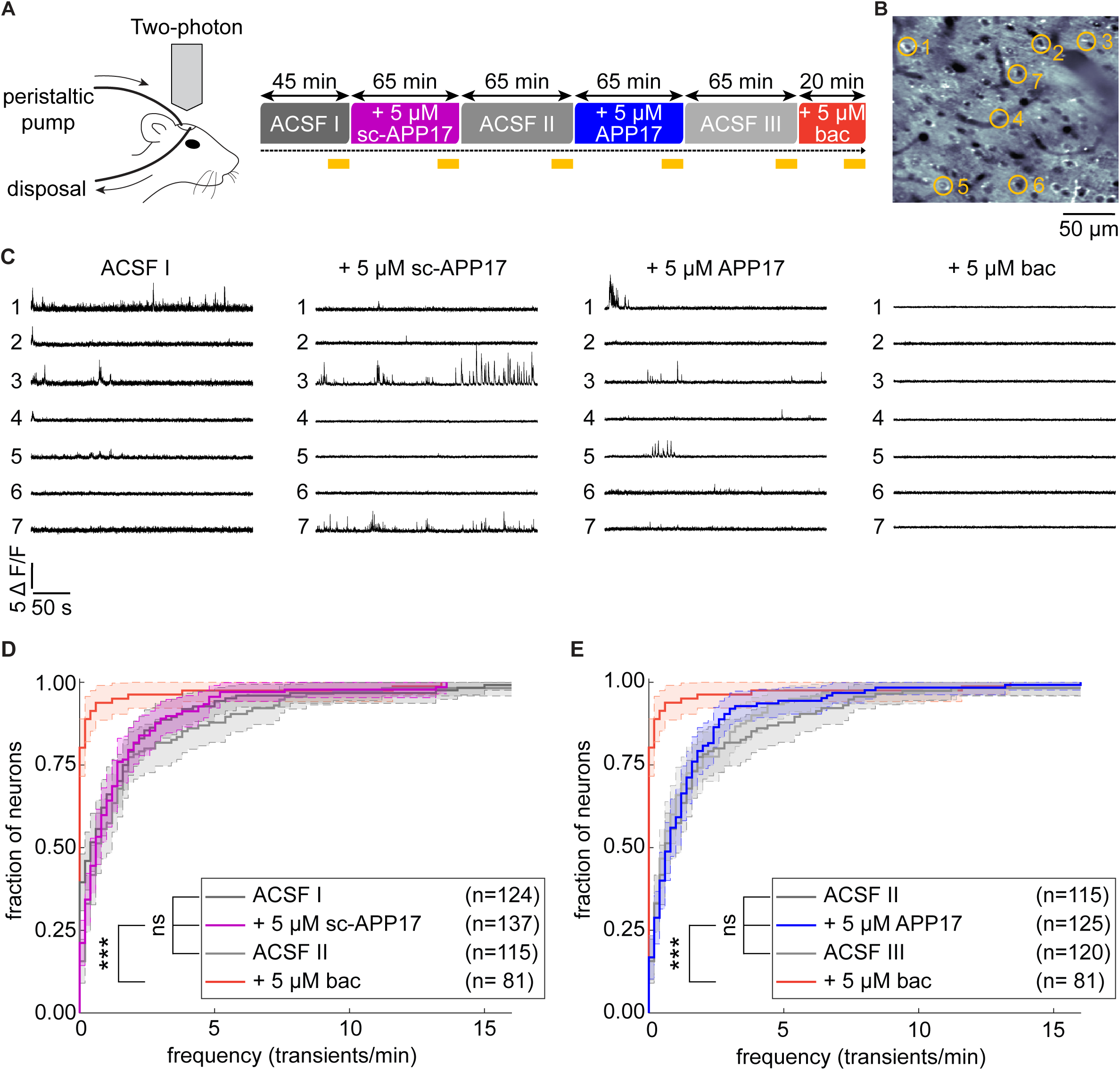
APP17 does not influence spontaneous neuronal activity in the auditory cortex of mice. (**A**) *Left* Two-photon imaging of Ca^2+^ transients in the auditory cortex of anesthetized mice during perfusion of ACSF, APP17, sc-APP17 and baclofen. *Right* Scheme of the experimental design. Time specifications denote the durations of the perfusions. Yellow lines indicate the two-photon Ca^2+^ imaging periods (5 min each). (**B**) *In vivo* two-photon image of neurons expressing GCaMP6f. Representative neurons selected to illustrate Ca^2+^ transients in (C) are marked with yellow circles. (**C**) Ca^2+^ transients of neurons shown in (B) across the entire 5 min imaging period of a given condition. (**D**) Cumulative distribution of the frequency of Ca^2+^ transients, comparing sc-APP17 with baseline (ACSF I) and washout (ACSF II) and perfusion with baclofen (bac). (**E**) Cumulative distribution of the frequency of Ca^2+^ transients, comparing APP17 with baseline (ACSF II) and washout (ACSF III) and baclofen. (D,E) 95% confidence intervals are shown as shaded areas. The number of neurons recorded in each condition are indicated. Kruskal Wallis multicomparison test: APP17 vs ACSF I, II, III and sc-APP17 are not significantly different (p > 0.05); bac vs ACSF I, II, III, sc-APP17 and APP17 are all significantly different (p < 0.0001). For detailed p-values, see Figure 9 – figure supplement 1. Source file containing Ca^2+^ transient data is available in Figure 9 – source data 1.

## Discussion

Proteolytic processing of APP through the non-amyloidogenic pathway liberates sAPP, which modulates synaptic functions, presumably by acting at neuronal cell surface receptors (Ishida et al., 1997; Bour et al., 2004; Taylor et al., 2008; Claasen et al., 2009; Aydin et al., 2011; Hick et al., 2015; Muller et al., 2017; Richter et al., 2018). Nanomolar concentrations of sAPP were shown to have PAM activity at heterologously expressed α7 nicotinic acetylcholine receptors, suggesting that nicotinic receptors mediate some of the effects of sAPP (Richter et al., 2018). Recent experiments identified GB1a/2 receptors as receptors for sAPP (Rice et al., 2019). GB1a/2 receptors are predominantly expressed at presynaptic sites, where they control neurotransmitter release (Vigot et al., 2006). It was shown that sAPP and APP17, a peptide of 17 amino acids corresponding to the SD1 binding-site of APP, reduce the frequency of mEPSCs and inhibit neuronal activity (Rice et al., 2019). While these findings received much attention and are consistent with activation of GBRs (Haass and Willem, 2019; Korte, 2019; Tang, 2019; Yates, 2019), fundamental questions remained. For example, it is unclear how a conformational change in SD1, induced by sAPP or APP17 binding, increases GBR activity. High-resolution structures of the GBR heterodimer in the apo, antagonist-bound, agonist-bound and agonist- and PAM-bound states in complex with the G protein are available and provide detailed insights into the activation mechanism of GBRs (Mao et al., 2020; Papasergi-Scott et al., 2020; Park et al., 2020; Shaye et al., 2020; Shaye et al., 2021; Shen et al., 2021). These structures show that the N-terminal SD1 is neither part of the binding sites for orthosteric or allosteric ligands, nor alters pharmacological receptor properties (Kaupmann et al., 1998) or participates in receptor activation (Evenseth et al., 2020; Shaye et al., 2021). Therefore, there is no straightforward explanation for potential functional effects of sAPP or APP17 at GBRs. Moreover, Rice and colleagues (Rice et al., 2019) did not analyze whether sAPP or APP17 regulate GB1a/2 receptors in heterologous cells, which is important to demonstrate a direct action at the receptor. In fact, in an earlier report showing interaction of native GBRs with APP, we found no evidence for recombinant sAPP protein regulating GB1a/2 receptors expressed in heterologous cells (Dinamarca et al., 2019). However, native GBRs form receptor complexes with additional proteins (Pin and Bettler, 2016; Schwenk et al., 2016; Bettler and Fakler, 2017) and these proteins could be necessary for the observed effects of APP17 on receptor activity.

The aim of this study was to clarify whether APP17 can activate recombinant and/or native GBRs. We could confirm that APP17 binds to purified recombinant SD1/2 protein, with a *K_D_* of 543 nM that is similar to the *K_D_* determined earlier (Rice et al., 2019). For functional experiments in HEK293T we used a range of established cell-based GBR assays reporting (1) conformational changes associated with receptor activation, (2) G protein activation, (2) cAMP inhibition and (4) Kir3 channel activation. In all these assays, APP17 had no agonistic, PAM or antagonistic properties at GB1/2 receptors. APP17 also did not influence constitutive GBR activity in the presence or absence of APP695 that competes with APP17 for binding at SD1. APP17 also did not modulate native GBRs in experiments assessing G protein activation in brain membranes, activation of K^+^ currents in cultured neurons, neurotransmitter release in acute hippocampal slices and neuronal activity in living mice. Thus, all our findings are consistent with a complete lack of functional effects of APP17 at GB1a/2 receptors, confirming the lack of functional effects observed with sAPP earlier (Dinamarca et al., 2019). The lack of functional effects is not due to a faulty APP17 peptide, since the APP17 peptide used in functional experiments was validated for binding recombinant SD1/2 protein and GB1a/2 receptors expressed in HEK293T cells. In all our experiments, we used GABA or baclofen to control for receptor activity. It therefore appears that sAPP mediates its neuronal effects through receptors other than GBRS. APP binding to GBRs probably mainly evolved to control receptor trafficking in axons and stabilize APP and GB1a/2 receptors at the cell surface (Hannan et al., 2012; Dinamarca et al., 2019). In principle, it is possible that sAPP interferes with the APP/GB1a interaction at the cell surface, which could lead to a downregulation of presynaptic GB1a/2 receptors and disinhibition of neurotransmitter release. However, the concentration of the abundant sAPPα variant in the interstitial fluid reaches ~ 1 nM (Dobrowolska et al., 2014). Considering a K*_D_* of 183 nM for the APP interaction with GB1a (Dinamarca et al., 2019), it is unlikely that endogenous levels of sAPP would reach concentrations high enough to displace APP from GB1a/2 receptors or to directly activate the receptor.

## Materials and methods

### Plasmids and reagents

The following plasmids were used: Flag-GB1a (Adelfinger et al., 2014); Flag-GB2, APP695 (Dinamarca et al., 2019); Gα_o_-RLuc, Venus-Gγ_2_ (Ayoub et al., 2009); Flag-Gβ_2_ (Rajalu et al., 2015); Myc-GB1a, HA-GB2 (Pagano et al., 2001); Kir3.1/Kir3.2 concatamer (Wischmeyer et al., 1997); PKA-Reg-RLuc-NT, PKA-Cat-RLuc-CT (Stefan et al., 2007) and SRE-FLuc (Cheng et al., 2010). GABA, CGP54626, forskolin, picrotoxin, and tetrodotoxin (TTX) were from Tocris Bioscience, Bristol, England.

### Peptide characterization

APP17 (Ac-DDSDVWWGGADTDYADG-NH_2_ (Rice et al., 2019)) and sc-APP17 (acetyl-DWGADTVSGDGYDAWDD-amide) peptides were from Insight Biotechnology, London, England (>98% purity). ESI-LC-MS (Poroshell, 300SB-C18, 2.1 × 75 mm, Agilent Technologies, Santa Clara, United States of America) and RP-UPLC (Acquity, Waters Corporation, Milford, United States of America) were used to confirm peptide mass and purity, respectively. ITC experiments were carried out in a microcalorimeter (Microcal ITC200, GE healthcare, Chicago, United States of America) at 25 °C with a stirring speed of 600 rpm in a buffer containing 20 mM NaPi (pH 6.8), 50 mM NaCl and 0.5 mM EDTA. For titration, APP17 or sc-APP17 (each 300 μM) were injected (first injection 0.5 μl, followed by 25 injections of 1.5 μl) into the sample cell containing purified recombinant SD1/2 protein (30 μM) (Schwenk et al., 2016). Control measurements of peptide versus buffer were subtracted from the peptide versus SD1/2 measurements. Data were analyzed with Microcal ITC200 Origin software, using a one-site binding model.

### Cell lines

Human Embryonic Kidney 293T (HEK293T) were directly obtained from ATCC (https://web.expasy.org/cellosaurus/CVCL_0063) and maintained in DMEM supplemented with 10% FBS (GE Healthcare) and 2% penicillin/streptomycin (Sigma-Aldrich, St. Louis, United States of America) at 37°C with 5% CO_2_. HEK293T cells stably expressing Gα_qi_ were a gift from the laboratory of Murim Choi (Seoul National University College of Medicine, Republic of Korea) (Yoo et al., 2017). All cell lines were authenticated using Short Tandem Repeat (STR) analysis by Microsynth (Switzerland) and tested negative for mycoplasma contamination.

### Cell culture and transfection

HEK293T cells were transiently transfected in Opti-MEM™ (Gibco, Thermo Fisher Scientific) using Lipofectamine™ 2000 (Thermo Fisher Scientific). The total amount of transfected DNA was kept equal by supplementing with pCI plasmid DNA (Promega, Madison, United States of America). For electrophysiological recordings, transfected cells were seeded on poly-L-lysine (Sigma-Aldrich) coated coverslips. Transfected cells were identified by their EGFP fluorescence. To establish primary cultures of hippocampal neurons, pregnant RjOrl:SWISS mice (Janvier Labs, France) were sacrificed under anesthesia by decapitation (Animal license number 1897_31476, approved by the Veterinary Office of Basel-Stadt, Switzerland). Dissected hippocampi of E17/18 embryos were collected in HBSS (Gibco, Thermo Fisher Scientific) and dissociated with 0.25% trypsin (Invitrogen, Thermo Fisher Scientific) at 37°C for 10 min. Cells were suspended in dissection medium [MEM Eagle (Sigma-Aldrich); 0.5% D(+)glucose; 10% horse serum (Gibco, Thermo Fisher Scientific); 0.1% Pen-Strep (Sigma-Aldrich)] to block trypsin activity. Cells were plated on 13 mm cell culture coverslips coated with 0.01 mg/ml poly-L-lysine hydrobromide (Sigma-Aldrich) at a density of 50,000 cells/cm^2^. Two hours after dissection, the medium was replaced with Neurobasal™ Medium (Gibco, Thermo Fisher Scientific) supplemented with B-27™ (Gibco, Thermo Fisher Scientific) and GlutaMAX™ (Thermo Fisher Scientific). Primary hippocampal neurons were maintained in a humidified incubator with 5% CO_2_ at 37 °C.

### APP17-TMR binding experiments

Transfected HEK293T cells expressing Flag-GB1a and Flag-GB2 were seeded into 96-well microplates (Greiner Bio-One, Kremsmünster, Austria) at 50,000 cells/well. After 18 hrs, peptides were mixed with conditioned medium at the following final concentrations: APP17-TMR (1 µM) with either APP17 (10 µM), sc-APP17 (10 µM) or PBS (vehicle); sc-APP17-TMR (1 µM) in PBS was used as a negative control. After removal of medium, peptide mixes were added to the wells and cells incubated in the dark for 1 hr at RT. After removal of the peptides, PBS with MgCl_2_ and CaCl_2_ (Sigma-Aldrich) was added to the wells. TMR fluorescence was monitored with a Spark® microplate reader (Tecan Group, Männerdorf, Switzerland) using a monochromator (Excitation 544 nM, 20 nM bandwidth; detection 594 nM, 25 nm bandwidth). TMR fluorescence was determined after subtraction of the sc-APP17-TMR fluorescence measured at HEK293T cells transfected with pCI plasmid.

### FRET measurements

Single and combined labelling of SNAP- and ACP-tag were performed as described previously (Lecat-Guillet et al., 2017). Briefly, 24 hrs after transfection, cells were incubated for 24 hrs at 30°C. The medium was removed and cells were incubated with 500 nM of SNAP-Red in Tag-Lite Buffer (Perkin Elmer Cisbio) for 1 hr at 37°C. Cells were washed once and incubated with 10 mM MgCl_2_, 1 mM DTT, 2µM CoA-Lumi4-Tb (Perkin Elmer Cisbio) and Sfp synthase (New England Biolabs) in Tag-Lite Buffer for 1 hr at 30°C. Cells were washed three times and APP17 or scAPP17 were added either alone or together with GABA in Tag-Lite Buffer. TR-FRET measurements were performed in Greiner black 96-well plates, using a PHERAstar FS microplate reader. After excitation with a laser at 337 nm (40 flashes per well), the fluorescence was collected at 620 nm (donor signal) and 665 nm (sensitized acceptor signal). The acceptor ratio was calculated using the sensitized acceptor signal integrated over the time window [50 µsec - 100 µsec], divided by the sensitized acceptor signal integrated over the time window [900 µsec - 1150 µsec].

### BRET measurements

BRET experiments were performed as described (Ivankova et al., 2013; Turecek et al., 2014; Dinamarca et al., 2019). HEK293T cells were transiently transfected with Flag-GB1a, Flag-GB2, Gα_o_-RLuc, Gβ_2_ and Venus-Gγ_2_ plasmids with or without APP695. In order to ensure APP695 binding to GB1a/2 *in trans* a pool of HEK293T cells expressing APP695 was mixed with a pool of HEK293T cells expressing Flag-GB1a, Flag-GB2, Gα_o_-RLuc, Gβ_2_ and Venus-Gγ_2_.Transfected cells were seeded into 96-well microplates (Greiner Bio-One) at 100,000 cells/well. After 18 hrs, cells were washed and coelenterazine h (5 µM, NanoLight Technologies, Prolume Ltd., Pinetop-Lakeside, United States of America) added for 5 min. Luminescence and fluorescence signals were alternatively recorded for a total of 845 sec using a Spark® microplate reader. Peptide, GABA or CGP54626 were injected with the Spark® microplate reader injection system at either 146 or 457 sec. The BRET ratio was calculated as the ratio of the light emitted by Venus-Gγ_2_ (530 – 570 nm) over the light emitted by Gα_o_-RLuc (370 – 470 nm). BRET ratios were adjusted by subtracting the ratios obtained when RLuc fusion proteins were expressed alone. Each data point represents a technical quadruplicate.

### PKA assay

PKA measurements were performed as described in (Stefan et al., 2007). HEK293T cells were transiently transfected with Flag-GB1a, Flag-GB2, PKA-Reg-RLuc-NT and PKA-Cat-RLuc-CT with or without APP695. Transfected cells were distributed into 96-well microplates (Greiner Bio-One) at a density of 80,000 cells/well. After 42 hrs, cells were washed and coelenterazine h (5 µM, NanoLight Technologies) added for 5 min. Luminescence signals were detected for a total of 1276 sec using a Spark® microplate reader. To induce PKA dissociation, 1 mM forskolin was added manually at 108 sec. Peptide, GABA or CGP54626 were injected at either 529 or 905 sec. Luminescence signals were adjusted to luminescence signals obtained by injecting PBS at 529 and 905 sec. The luminescence was normalized to baseline luminescence. Curves were plotted after forskolin addition and the time point 71 sec prior the first injection was set to 0. Each data point represents a technical quadruplicate.

### SRE-luciferase accumulation assay

HEK293T cells stably expressing Gα_qi_ were transiently transfected with Flag-GB1a, Flag-GB2 and SRE-FLuc with or without APP695. In order to ensure GB1a/2 binding of APP695 *in trans* a pool of HEK293-Gα_qi_ cells expressing APP695 was mixed with HEK293-Gα_qi_ cells expressing Flag-GB1a, Flag-GB2 and SRE-FLuc. Transfected cells were distributed into 96-well microplates (Greiner Bio-One) at a density of 80,000 cells/well. After 24 hr, the culture medium was replaced with Opti-MEM™-GlutaMAX™. Peptides were incubated in Opti-MEM™-GlutaMAX™ for 1 hr. In presence of peptide, GB1a/2 receptors were activated with various concentrations of GABA for 15 hr. FLuc activity in lysed cells was measured using the Luciferase® Assay Kit (Promega) using a Spark® microplate reader. Luminescence signals were adjusted by subtracting the luminescence obtained when expressing SRE-FLuc fusion proteins alone.

### Electrophysiology

Neuronal cultures and HEK293T cells were prepared as described (Dinamarca et al., 2019). Coverslips with hippocampal neurons (DIV 12-15) or HEK293T cells were transferred to a chamber containing a low-K^+^ bath solution (in mM): 145 NaCl, 4 KCl, 5 HEPES, 5.5 D-glucose, 1 MgCl_2_ and 1.8 CaCl_2_ (pH 7.4 adjusted with NaOH). Recordings were performed at room temperature using borosilicate pipettes of 3-5 MΩ resistance tips, filled with K-gluconate-based pipette solution (in mM): 150 K-gluconate, 1.1 EGTA, 10 HEPES, 10 Tris-phosphocreatine, 0.3 NaGTP and 4 MgATP (pH 7.2 adjusted with KOH). Upon achieving whole-cell access, cells were held in voltage-clamp mode at −70 mV (with no correction for liquid junction potential) and baclofen-induced K^+^ currents were induced in a high-K^+^ bath solution (in mM): 120 NaCl, 25 KCl, 5 HEPES, 5.5 D-glucose, 1 MgCl_2_ and 1.8 CaCl_2_ (pH 7.4 adjusted with NaOH). Whole-cell patch-clamp recordings from visually identified CA1 pyramidal cells in acute hippocampal slices of juvenile male and female C57BL/6JRj mice (Janvier Labs, France) were performed as described (Vigot et al., 2006) (animal license number 1897_31476, approved by the Veterinary Office of Basel-Stadt, Switzerland). Schaffer collaterals were stimulated at 0.1 Hz with brief current pulses via bipolar Pt/Ir wires. Evoked EPSCs were recorded at −60 mV with a Cs^+^-based intracellular solution. Baclofen and peptides were bath applied in standard ACSF at room temperature.

### [^35^S]GTPyS binding

Preparation of mouse brain membranes was performed as described earlier (Olpe et al., 1990). Briefly, 8 weeks old male C57BL/6JRj mice (Janvier Labs, France) were decapitated under isoflurane anesthesia (animal license number 1897_31476, approved by the Veterinary Office of Basel-Stadt, Switzerland). The brains were removed, washed in ice-cold PBS and homogenized in 10 volumes of ice-cold 0.32 M sucrose, containing 4 mM HEPES, 1 mM EDTA and 1 mM EGTA, using a glass-teflon homogenizer. Debris was removed at 1,000 *g* for 10 min and membranes were centrifuged at 26,000 *g* for 15 min. The pellet was osmotically shocked by re-suspension in a 10-fold volume of ice-cold H_2_O and kept on ice for 1 hr. The suspension was centrifuged at 38,000 *g* for 20 min and re-suspended in a 3-fold volume of H_2_O. Aliquots were frozen in liquid nitrogen and stored at −20°C for 48 hrs. After thawing at room temperature, a 7-fold volume of Krebs-Henseleit (KH) buffer (pH 7.4) was added, containing 20 mM Tris-HCl, 118 mM NaCl, 5.6 mM glucose, 4.7 mM KCl, 1.8 mM CaCl_2_, 1.2 mM KH_2_PO_4_ and 1.2 mM MgSO_4_. Membranes were washed three times by centrifugation at 26,000 *g* for 15 min, followed by re-suspension in KH buffer. The final pellet was re-suspended in a 5-fold volume of KH buffer. Aliquots of 2 ml were frozen and stored at −80 °C until the day of the experiment. On the day of the experiment, frozen membranes were thawed, homogenized in 10 ml ice-cold assay buffer I containing 50 mM Tris-HCl buffer (pH 7.7); 10 mM MgCl_2_, 1.8 mM CaCl_2_, 100 mM NaCl, and centrifuged at 20,000 *g* for 15 min. The pellet was re-suspended in the same volume of cold buffer and centrifuged twice as above with 30 min of incubation on ice in between the centrifugation steps. The resulting pellet was re-suspended in 150 μl of assay buffer II (per point) containing 50 mM Tris-HCl buffer (pH 7.7); 10 mM MgCl_2_, 1.8 mM CaCl_2_, 100 mM NaCl, 30 μM guanosine 5‘-diphosphate (GDP; Sigma-Aldrich) and 20 μg of total membrane protein. To this, 8 μM of the APP17 peptide was added in 25 μl of phosphate buffer (50 mM sodium phosphate, pH 6.8, 50 mM NaCl, for + APP17) or 25 μl of phosphate buffer alone (for – APP17) and incubated for 30 min. The reaction was started by adding various concentrations of GABA and 0.2 nM of [^35^S]GTPγS (PerkinElmer, Waltham, United States of America) in a final volume of 200 μl per point and assayed as described earlier (Rajalu et al., 2015). Non-specific binding was measured in the presence of unlabeled GTPγS (10 μM; Sigma-Aldrich). The reagents were incubated for 1 hr at room temperature in 96-well polypropylene microplates (Greiner Bio-One) with mild shaking. They were subsequently filtered using 96-well Whatman GF/C glass fiber filters (PerkinElmer), pre-soaked in assay buffer, using a Filtermate cell harvester (PerkinElmer). After four washes with assay buffer, the Whatman filter fibers were dried for 2 hrs at 50 °C. 50 μl of scintillation fluid (MicroScint™-20; PerkinElmer) was added, the plates were shaken for 1 hr and thereafter counted using a Packard TopCount NXT (PerkinElmer).

### In vivo two-photon Ca^2+^ imaging of auditory cortex

Ca^2+^ imaging experiments were approved by the Veterinary Office of Basel-Stadt, Switzerland (animal license number 3004_34045). We crossed Ai95(RCL-GCaMP6f)-D mice (Madisen et al., 2015) (RRID:IMSR_JAX:028865) with Nex-Cre mice (Goebbels et al., 2006) to obtain GCaMP6f expression in cortical neurons. 9-12 weeks old male mice were anesthetized with isoflurane (4% induction, 1.5–2.5% surgery, 1% optical imaging). 3.2 mg/kg dexamethasone was administered intraperitoneally 48, 24 and 1 hr prior to surgery to prevent brain swelling. Bupivacaine/lidocaine (0.01/0.04 mg) was injected subcutaneously for analgesia. An imaging chamber was created with cement above right auditory cortex (rACx) and a post fixed on the skull of the left hemisphere. A craniotomy (1.5 x 1.5mm^2^) above rACx was carefully opened and duratomy was performed. For optical imaging, the post was stably fixed on a stage and the head tilted 30° for optimal access to the rACx. The imaging chamber was perfused at 1ml / min using a peristaltic pump. All solutions were kept at 37°C for 1 hr prior to perfusion. Two-photon Ca^2+^ imaging periods (5 min each) started 45 min after the first ACSF perfusion, 60 min after sc-APP17, second ACSF, APP17 and third ACSF perfusion and 15 min after baclofen perfusion. Ca^2+^ transients of neuronal cell bodies in the upper layers of rACx (focal depth: 150 - 250 µm) were recorded with a two-photon microscope (INSS) equipped with an 8 kHz resonant scanner. Images were acquired with a PXI-1073-based data acquisition system (NI) through a Nikon 16x objective (0.8 NA), at 30Hz within a 500 x 500 μm field of view (512 x 512 pixels). The wavelength to excite GCaMP6f was 940 nm (Chameleon Vision-S, Coherent). We used the optical imaging control and data acquisition software ScanImage 5.7 (Pologruto et al., 2003). Correction of Z-drift and motion artifacts, detection of neuronal cell bodies and extraction of Ca^2+^ signals was with the Python 3 based image processing pipeline suite2p (Pachitariu et al., 2017). Data analysis visualization and statistics were performed using custom-written MATLAB scripts (Mathworks, Natick, MA). Automated detection of Ca^2+^ transients was with an adapted algorithm (Sorensen et al., 2017). First, the fluorescence signal was corrected by the median filtered data removing slow trends. The detection threshold was set to 2.5 times the standard deviation and a minimum peak width of 3 data points above threshold to remove fast artifacts. ΔF/F was calculated as (F-F_0_)/F_0_, were F_0_ is the mean fluorescence across all detected neurons in a given condition. The rates of Ca^2+^ transients representing the recorded neuronal populations for each condition were plotted as the empirical cumulative distribution function with 95% confidence intervals.

## Statistical analysis

Data was analyzed with GraphPad Prism version 8 (GraphPad, San Diego, United States of America) if not indicated otherwise. Sample size in all experiments was based on those of similar experiments in previous studies. Samples were randomly allocated to the experimental groups. Confounding effects in the cell-based assays (Figs. 2-5) were minimized by rotating the order of cell plating. Blinding was not performed for any experiment. Individual data sets were tested for normality with the Shapiro-Wilk or D’Agostino-Pearson test (for n ≥ 8). For the datasets obtained with the cell-based assays (Figs. 2-5) outliers were identified using the ROUT method with Q = 1%. For all other experiments, no data was excluded. Statistical significance of data sets against 0 or 100 was assessed by one sample t-test. For non-normal distribution, the non-parametric one sample Wilcoxon test was used. Statistical significance between two groups containing one variable was assessed by student’s t-test. Statistical significance between three or more groups containing one variable was assessed by ordinary or paired one-way ANOVA with Holm-Sidak’s multiple comparisons test. For non-normal distribution, the non-parametric Friedman test with Dunn’s multiple comparisons test was used. Statistical significance between groups containing two variables was assessed by ordinary two-way ANOVA with Sidak’s multiple comparisons test. Statistical significance between dose-response curves was assessed by extra sum-of-squares F test of non-linear regression curve fits. P-values < 0.05 were considered significant. Data are presented as mean ± standard error of mean (SEM) or mean ± standard deviation (SD) as indicated in the figure legends.

## Data availability

For all figures, numerical data that are represented in graphs are provided as source data excel files.

## Acknowledgments

The Swiss National Science Foundation (31003A-152970 to B.B.) supported this work. P.D.R., P.R., J.-P.P., J.S., B.F., M.G., T.R.B., K.S. and B.B. conceived the project. P.D.R., V.S., S.R., D.F.-F., D U., T.F., L T., S.R., Z.C., and M.G. performed experiments. P.D.R. and B.B. wrote the manuscript with the help of the other authors. B.B. is a member of the scientific advisory board of Addex Therapeutics, Geneva. K.S. is a co-founder and a part-time employee of Avilex Pharma. All other authors declare no conflict of interest.

## Legends for figure supplements

**Figure 5 – figure supplement 1.**
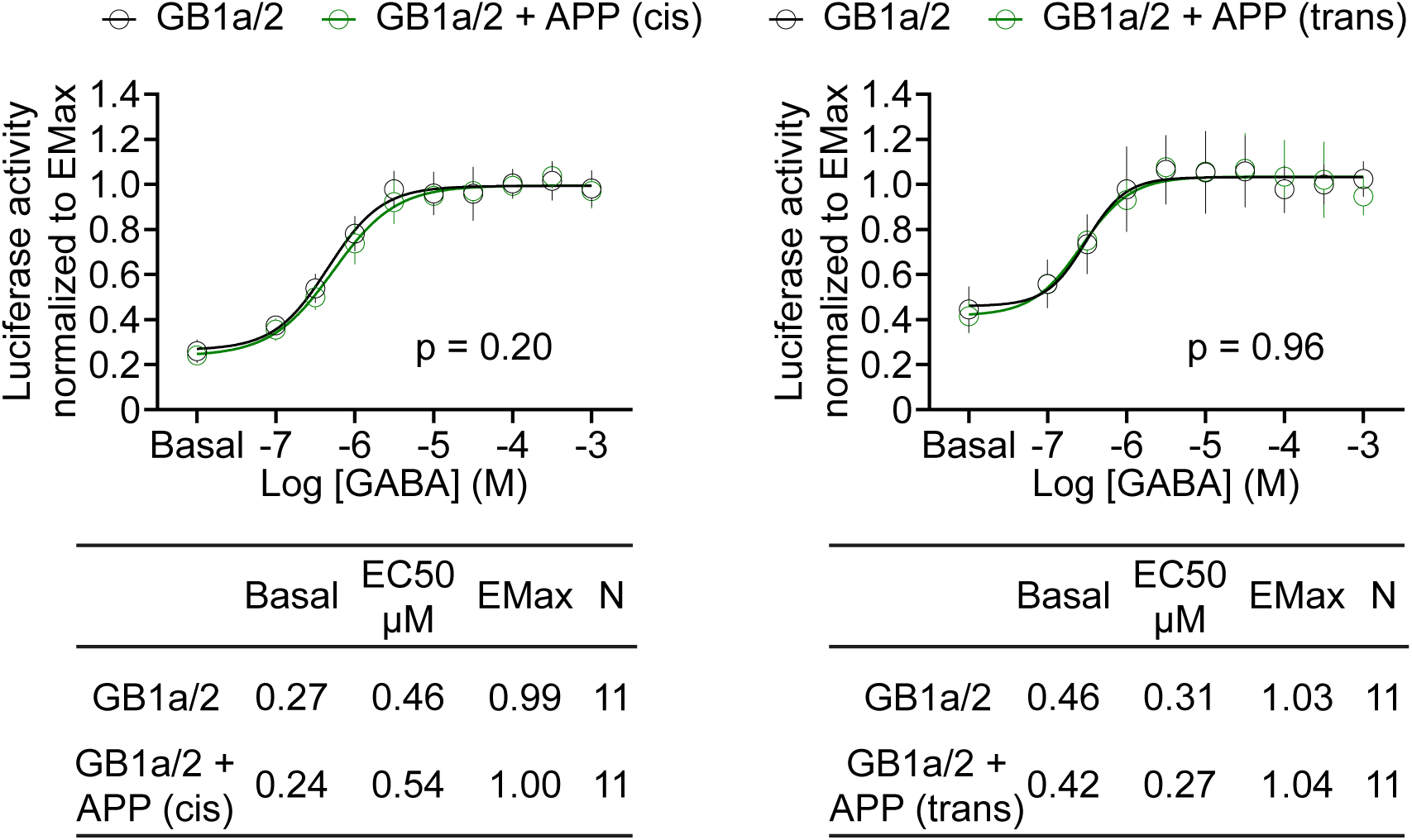
APP695 expressed *in cis* or *in trans* with GB1a/2 receptors exerts no allosteric effects on Gα_qi_ signaling in HEK293T cells. GABA dose-response curves show no difference in the absence (black) or presence of APP695 (green) *in cis* (left) or *in trans* (right). Tables show basal, EC_50_ and Emax values derived from the curve fits. All data are means ± SD. The number of independent experiments is indicated in the tables. Non-linear regression curve fits of 11 independent experiments per condition. p = 0.20, p = 0.96, extra sum-of-squares F test. Source file containing FLuc activity data is available in Figure 5 – source data 1.

**Figure 7 – figure supplement 1.**
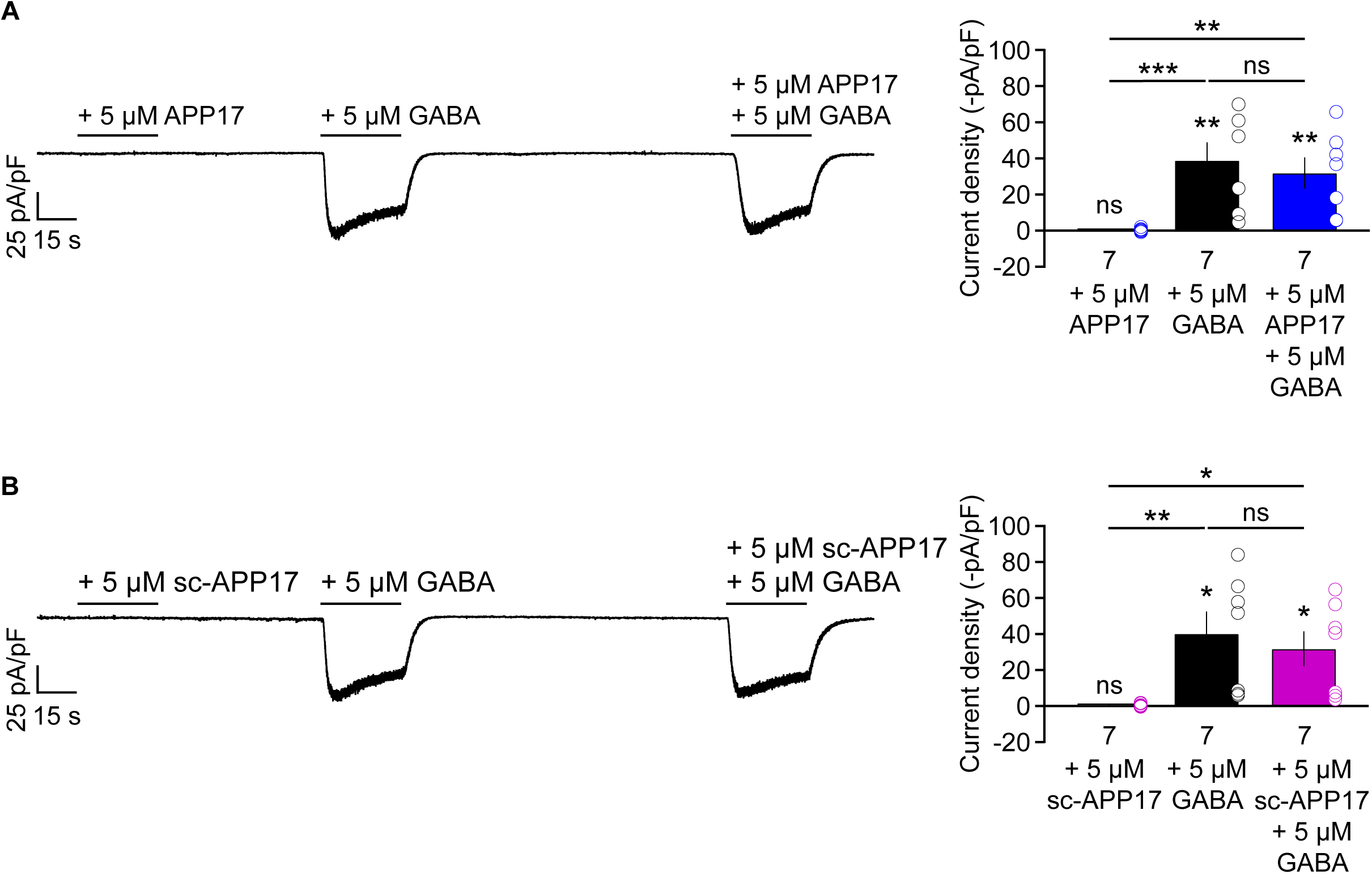
APP17 does not evoke or influence GB1a/2 receptor-induced Kir3 currents in transfected HEK293T cells. (**A**,**B**) *Left* Representative traces showing that neither APP17 (A) nor sc-APP17 (B) evoke GB1a/2 receptor-induced K^+^ currents in transfected HEK293T cells. Application of GABA alone or in the presence of APP17 (A) or sc-APP17 (B) yielded similar current amplitudes, showing that the peptides do not allosterically modulate GABA-induced currents. *Right* Bar graphs showing K^+^ current densities determined in experiments as shown to the left. Data are means ± SEM. The number of independent experiments is indicated in the bar graphs. ns = not significant, *p < 0.05, **p < 0.01, ***p < 0.001 Paired one-way ANOVA with Holm-Sidak’s multiple comparisons test (to compare different means) and One sample t-test against 0. Source file containing K^+^ current data is available in Figure 7 – source data 1.

**Figure 9 – figure supplement 1.**
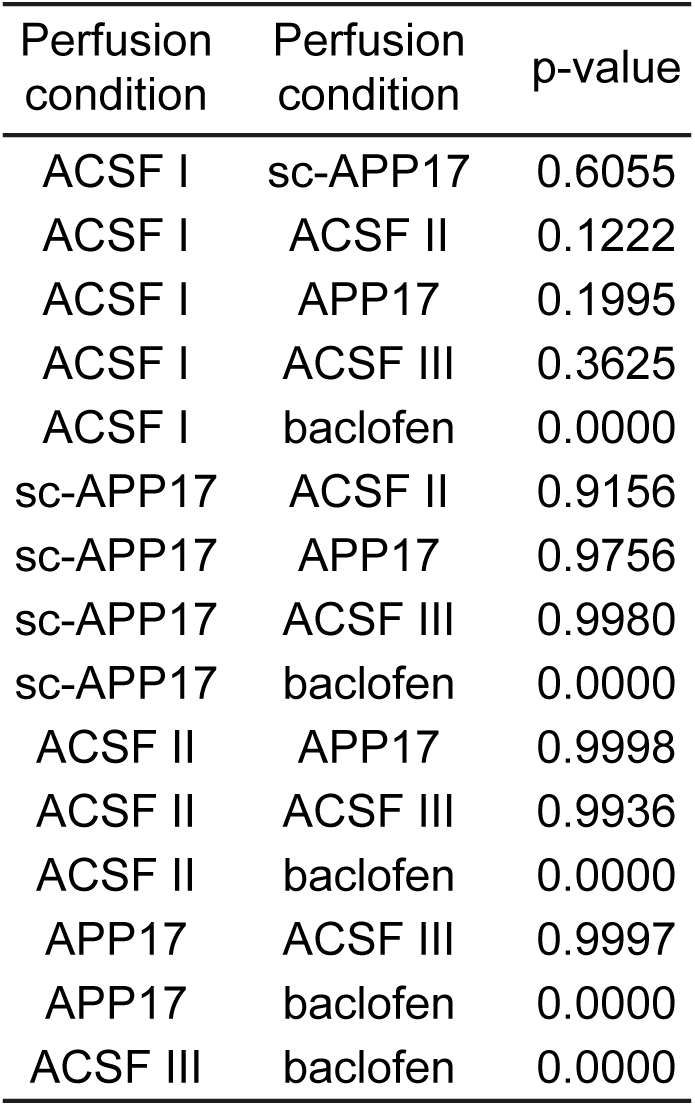
Statistical analysis between perfusion conditions in two-photon Ca^2+^ imaging experiments. The significance between experimental conditions was tested with the Kruskal-Wallis multicomparison test as the data was not normally distributed. Note, only baclofen is significantly different (p < 0.0001) from all other experimental conditions.

